# Characterising the mechanism of action of an ancient antimicrobial, honey, using modern transcriptomics

**DOI:** 10.1101/2020.02.12.946830

**Authors:** Daniel Bouzo, Nural N. Cokcetin, Liping Li, Giulia Ballerin, Amy L. Bottomley, James Lazenby, Cynthia B. Whitchurch, Ian T. Paulsen, Karl A. Hassan, Elizabeth J. Harry

## Abstract

Manuka honey has broad-spectrum antimicrobial activity and unlike traditional antibiotics, resistance to its killing effects has not been reported. However, its mechanism of action remains unclear. Here we investigated the mechanism of action of manuka honey and its key antibacterial components using a transcriptomic approach in a model organism, *Pseudomonas aeruginosa.* We show that no single component of honey can account for its total antimicrobial action, and that honey affects the expression of genes in the SOS response, oxidative damage and quorum sensing. Manuka honey uniquely affects genes involved in the explosive cell lysis process and in maintaining the electron transport chain, causing protons to leak across membranes and collapsing the proton motive force; and induces membrane depolarisation and permeabilisation in *P. aeruginosa*. These data indicate that the activity of manuka honey comes from multiple mechanisms of action that do not engender bacterial resistance.

**Importance:** The threat of antimicrobial resistance to human health has prompted interest in complex, natural products with antimicrobial activity. Honey has been an effective topical wound treatment throughout history, predominantly due to its broad-spectrum antimicrobial activity. Unlike traditional antibiotics, honey-resistant bacteria have not been reported, however, honey remains underutilised in the clinic in part due to a lack of understanding of its mechanism of action. Here we demonstrate that honey affects multiple processes in bacteria, and this is not explained by its major antibacterial components. Honey also uniquely affects bacterial membranes and this can be exploited for combination therapy with antibiotics that are otherwise ineffective on their own. We argue that honey should be included as part of the current array of wound treatments due to its effective antibacterial activity that does not promote resistance in bacteria.

## Introduction

Honey has been used for millennia as a topical antibacterial ^1–6^ and, unlike traditional antibiotics, bacterial resistance to honey has not been reported ^7, 8^. The increasing prevalence of antimicrobial resistance demands alternative infection control and has prompted renewed scientific interest in complex, natural products with potent antimicrobial activity, like honey. However, honey remains underutilised in the clinic presumably due to a paucity of information on the mechanisms by which honey kills bacteria.

Honey is a complex mixture, with over 100 components, including sugars, proteins, phenols and plant- and bee-derived enzymes ^9^. The antibacterial activity of honey is derived from multiple factors: osmotic stress from the high sugar concentration ^10, 11^; low pH (between 3.2– 4.5); and, the presence of hydrogen peroxide (H_2_O_2_) produced from the bee-derived enzyme glucose oxidase. It was widely considered that the latter was the primary source of the antibacterial activity of honey and it is known to vary significantly in honeys from different floral sources ^10, 12–15^, however, following neutralisation of H_2_O_2_ by catalase, certain honeys retained high levels of antibacterial activity, referred to as non-peroxide activity (NPA). NPA was first observed in New Zealand manuka (*Leptospermum scoparium*) honey ^15, 16^. It has now been established that active manuka-type (*Leptospermum* sp.) honeys from New Zealand and Australia have substantially higher levels of NPA compared to honeys from other floral sources ^17, 18^. This is, in part, due to the high concentrations of the naturally occurring chemical methylglyoxal (MGO) in some *Leptospermum-*derived honeys ^19, 20^.

While MGO is a key antibacterial component of manuka honey, it alone cannot account for its total antimicrobial activity ^21–23^, as manuka honey inhibits the growth of pathogenic bacteria (including *Pseudomonas aeruginosa*, *Escherichia coli* and *Staphylococcus aureus*) at concentrations well below the minimum inhibitory concentration (MIC) of MGO alone ^21–24^. Additionally, many bacteria are innately equipped to detoxify MGO ^25–27^, so additional components in honey must also modulate its activity. From this, we hypothesise that the antibacterial activity of manuka honey comes from a combination of its various constituents and that its mechanism of action cannot be elucidated based exclusively on investigations of the individual components. Rather, to generate a fundamental understanding of the mechanism of antibacterial activity, the effects of the key components of manuka honey against microorganisms must be studied in isolation and in combination with each other. Despite the prominent role of MGO in the antibacterial activity of manuka honey, to what degree it contributes to the effect manuka honey has on bacterial gene expression and physiology has not been thoroughly investigated ^28–34^. Currently, the antimicrobial activity of manuka honey is reported and marketed based on its NPA, which can be directly tested via bioassays or derived from the MGO concentrations of manuka honey since MGO and NPA are well correlated^18^. This is problematic since NPA is only a measure of anti-staphylococcal activity and not representative of activity against other bacterial species ^35^. Therefore, it is important to understand how MGO alone and in combination with sugars works against Gram-negative microorganisms like *P. aeruginosa*, in order to better understand the mechanism of whole manuka honey. This is critical for its use in infection control, which requires killing of multiple species of bacteria present in wounds.

Previous studies have identified a number of biological processes in bacteria that may be affected by the action of honey, including cell division ^22, 29, 32, 33^, motility ^28^, quorum sensing (QS) ^36–40^, protein synthesis ^29, 32, 41^ and responses to oxidative stress ^7, 41^. With the increased affordability, sensitivity and accessibility of genetic analysis, we can now elucidate the entire changes that happen to a bacterial cell when exposed to different treatments. We have used a global transcriptomic approach, RNA-seq, as well as classical cell biology techniques, to characterise the effects of manuka honey and its key components (MGO, sugar and the combination) on *P. aeruginosa*. This opportunistic pathogen is commonly associated with burn wounds and surgical site infections ^42^ and listed by the World Health Organisation as a Priority 1 ‘critical’ pathogen for which novel treatment therapies are urgently needed ^43^. We demonstrate that 1) exposure to manuka honey causes significant, widespread changes in the transcriptomic profile of *P. aeruginosa*; 2) the mechanism of action and effect of honey on the transcriptomic response of *P. aeruginosa* is different to that of MGO, sugar or a combination thereof; and 3) only whole manuka honey, and not these key components, dissipates membrane potential in *P. aeruginosa* and is an important part of the mode of action of manuka honey.

## Methods

### Bacterial strains, media and antimicrobial agents

The bacterial strains used in this study are described in Table S1. Strains were cultured on cation-adjusted Mueller Hinton (CAMH media grown aerobically at 37°C unless stated otherwise.

**Table 1.** MIC and MBC values of manuka honey and honey analogues against P. aeruginosa PA14 ^a^. Quantity of MGO in parts per million (ppm) present at MIC concentrations of each sample.

| Sample | MIC (% w/v ± SD) | MBC (% w/v ± SD) | MGO content at<br>MIC (ppm) |
| --- | --- | --- | --- |
| Manuka honey | 10 ± 0.25 | 12 ± 0.25 | 90 |
| Artificial honey | 25 ± 0.5 | > 30 ± 0.00 | 0 |
| MGO | 55 ± 0.00 | > 55 ± 0.00 | 495 |
| Artificial honey + MGO | 21.5 ± 0.5 | > 30 ± 0.00 | 193.5 |

<sup>a</sup> MICs and MBCs expressed as the mean percentage (w/v) ± SD from three separate trials all performed in triplicate.
Percent concentrations for manuka honey components are presented as relative to their concentration within whole manuka.

The manuka honey used in this study is the same unprocessed honey, collected and prepared as previously described ^23^ (MGO: 958 mg/kg; H_2_O_2_: 0.34 µmol/h) and was supplied by Comvita Ltd, New Zealand. Artificial honey (AH) was made as sugar solutions of fructose (41.4 %), glucose (37.3%), sucrose (2.9%) and water (18.4%) (w/v) ^44^, and served as a means to measure the effects of the sugar component of honey. Methylglyoxal (Sigma-Aldrich) only treatment (MGO) was prepared as an aqueous solution at a final concentration equivalent to the manuka honey sample (958 mg/kg). This served as a measure of the contribution of MGO relative to the manuka honey. An artificial honey doped with methylglyoxal (AHMGO) was prepared as per the AH recipe above, with the modification of adding MGO, at a concentration equivalent to the manuka honey sample (958 mg/kg), to the water component prior to solubilising the sugars. AH, AHMGO and MGO samples were adjusted to pH 4.6 (the native pH of manuka honey) using sodium citrate then filter sterilised ^44^. All samples were stored in the dark at 4°C and freshly diluted prior to each experiment. Concentrations are reported as % weight per volume (w/v) in this study.

### Determination of Minimal Inhibitory Concentration (MIC), Minimum Bactericidal Concentration (MBC) and synergistic interaction of antimicrobial agents

MICs of all treatments were determined using the broth microdilution method previously described ^45^ with minor changes. CAMH broth was used for all assays and the final concentration of inoculum was 5×10^5^ CFU/mL. MBCs were determined by inoculating fresh CAMH agar plates from wells ≥ MIC of MIC plates with a sterile wooden stick and checking for growth after 24 hour incubation at 37°C. For susceptibility testing under anaerobic conditions, cultures were grown in CAMH broth supplemented with 1% KNO_3_ (Sigma-Aldrich) and anaerobic conditions were achieved using an Anoxomat^®^ II system (Mart Microbiology BV). Antimicrobial interactions with honey were characterised by standard chequerboard as previously described ^46^, however the final inoculum concentration was 5×10^5^ CFU/mL. Synergistic, antagonistic and no interactions were determined using the Fractional Inhibitory Concentration Index (FICI) ^47^.

### Total RNA isolation

*P. aeruginosa* PA14 cultures were prepared in CAMH broth to an initial OD_600_ of 0.05 and then incubated in a 250 mL cell culture flask (Falcon, Corning) at 37°C with shaking at 200 rpm until reaching mid-exponential phase (OD_600_ 0.4). Cultures were then split into four flasks containing 2 mL of each treatment at a final concentration of 0.5× MIC; manuka honey (5% w/v), artificial honey (12.5% w/v), artificial honey with MGO (10.75% w/v) and MGO solution (27.5% w/v). A fifth flask containing 2 mL of fresh CAMHB remained untreated. Cultures were grown for an additional 30 minutes before and RNA extraction was carried out as previously described ^48^. Experiments were conducted in triplicate and samples sent to Macrogen (Seoul, South Korea) for Ribosomal RNA reduction using Ribo-Zero^™□^ rRNA Removal kit (Illumina), library preparation using the TruSeq^Ⓡ^ stranded mRNA kit (Illumina) and subsequent 100 bp paired-end RNA sequencing on a HiSeq4000 sequencer (Illumina).

### Bioinformatic analysis

RNA-seq read quality was assessed using FASTQC (version 0.11.5) and trimmed using Trimmomatic (version 0.36) using the default parameters and trimmed of adaptor sequences (TruSeq3 paired-ended). Reads were aligned to the *P. aeruginosa* UCBPP-PA14 genome (http://bacteria.ensembl.org/Pseudomonas_aeruginosa_ucbpp_pa14/Info/Index/, assembly ASM14162v1) and then counted using the RSubread aligner (version 1.30.7) with the default parameters ^49^. After mapping, differential expression analysis was carried out using strand-specific gene-wise quantification using the DESeq2 package (version 1.18.0) ^50^. Further normalisation was conducted using RUVSeq (version 1.13.0) and the RUV correction method with *k = 1*, to correct for batch effects, using replicate samples to estimate the factors of unwanted variation ^51^. Absolute counts were transformed into standard z-scores for each gene over all treatments – that is absolute read for a gene minus mean read count for that gene over all samples and then divided by the standard deviation of all counts over all samples. Genes with an adjusted *P* value (p.adj) ≤ 0.05 were considered differentially expressed. PseudoCAP analysis was conducted by calculating the percentage of genes in each classification that were differentially expression (log_2_FC ≥ ±2, p.adj ≤ 0.05). Classifications were downloaded from the *Pseudomonas* Community Annotation Project ^52^.

### Assessment of membrane potential after antimicrobial treatment

Liposomes were formed using *Escherichia coli* polar lipid extract (Avanti Polar Lipids). Lipids were dried under argon or nitrogen from a chloroform suspension to form a lipid film in a glass tube. The lipids were suspended in liposome buffer (25 mM HEPES-NaOH (pH 7.0), 200 mM NaCl, 1 mM dithiothreitol) and subjected to 11 passages of extrusion each through 0.4 μm then 0.2 μm polycarbonate. Five hundred microliter samples were prepared in the same buffer including 5 mg of preformed liposomes, 1 mM pyranine (8-hydroxypyrene-1,3,6-trisulfonic acid trisodium salt) and 1.1 % *n*-octylglucoside. The samples were incubated at room temperature for 15 minutes, then diluted 1:60 with cold liposome buffer to dilute the *n*-octylglucoside to a concentration below its critical micelle concentration. The diluted samples were ultracentrifuged (185,000 × *g*) for 2 hours to collect the liposomes that were resuspended in 100 μL of liposome buffer. For each experiment, liposomes were diluted 1:100 into assay buffer (25 mM HEPES-KOH (pH 7.0), 200 mM KCl), and pyranine fluorescence was continuously monitored to detect pH changes in the lumen of the liposomes [F_509_ (em 450 nm)/ F_509_ (em 400 nm)]. A low concentration (5 nM) of the potassium ionophore valinomycin was added to facilitate the formation of an electrical gradient across the membrane, followed by whole manuka honey, or honey components. In control experiments, the polarity of the electrical gradient was reversed by reversing the isosmolar sodium and potassium salts in the liposome and assay buffers.

Membrane potential was assessed using the voltage-sensitive fluorophore DiBAC_4_(3) and the membrane-impermeable dye TO-PRO^®^-3. Mid-exponential phase cells (OD_600_ 0.4) were treated with 10% w/v of either manuka honey, AH, AHMGO, MGO or the positive control of 100 µM CCCP for 120 minutes at 37°C with shaking at 200 RPM. Then 10 µL of each sample was added to 490 µL of PBS containing 0.1 nM TO-PRO^®^-3 (Life Technologies) and 0.5 µM DiBAC_4_(3) (Sigma-Aldrich) (final DMSO concentration did not exceed 1% v/v) and left to incubate at room temperature in the dark for 15 minutes. Samples were then analysed on a BD LSRII flow cytometer for forward scatter (FSC), side scatter (SSC), fluorescein isothiocyanate (FITC)-A and allophycocyanin (APC) fluorescence. A total of 30,000 events were recorded for each sample and gated based on FSC and SSC. Subsequent analysis of flow cytometry data was conducted using FlowJo software (version 10.5.0).

## Results and discussion

### The antimicrobial activity of manuka honey against *P. aeruginosa* cannot be explained solely by methyglyoxal presence or levels

We determined the contribution of MGO and sugar (either in isolation or in combination with one another) to the antibacterial activity of manuka honey against *P. aeruginosa* by minimum inhibitory and minimum bactericidal concentration (MIC and MBC, respectively) assays (Table 1).

*P. aeruginosa* growth was inhibited by 10% w/v manuka honey, in agreement with previous reports ^53^ and manuka honey was bactericidal at 12% w/v (Table 1). The MIC values for AH, AHMGO and MGO alone were higher than that of manuka honey and MGO had the highest MIC of all treatments; none of these treatments were bactericidal at the highest concentrations that could practicably be tested (Table 1). The MIC for MGO alone was 5.5-fold higher than that of manuka honey (equivalent to 495 ppm MGO compared to 90 ppm MGO, respectively) similar to previous reports for the MIC of MGO against *P. aeruginosa* PAO1 ^24^. These results show that although MGO does contribute to the activity of manuka honey against *P. aeruginosa*, it is not the main factor responsible for inhibition or cell death. This is very different to *S. aureus* where the MGO content of manuka honey correlates strongly with anti-staphylococcal activity ^18^.

### Manuka honey affects a range of biological processes and pathways

The molecular responses of *P. aeruginosa* to manuka honey and its key components was investigated using RNA-Seq. We applied treatments at subinhibitory concentrations and short exposure times (0.5× MIC for 30 minutes), since this approach induces more specific responses and reduces indirect effects, thereby giving the most informative transcriptomic data ^54, 55^. We confirmed these conditions induced significant and meaningful changes in gene expression by pilot RT-qPCR experiments on targeted genes (Figure S1). It should be noted that using 0.5× MIC across all treatments meant that the final MGO concentration was different under each treatment condition in the RNA-Seq experiments (Table S2), and we were mindful of this in interpreting the data.

**Table 2.**
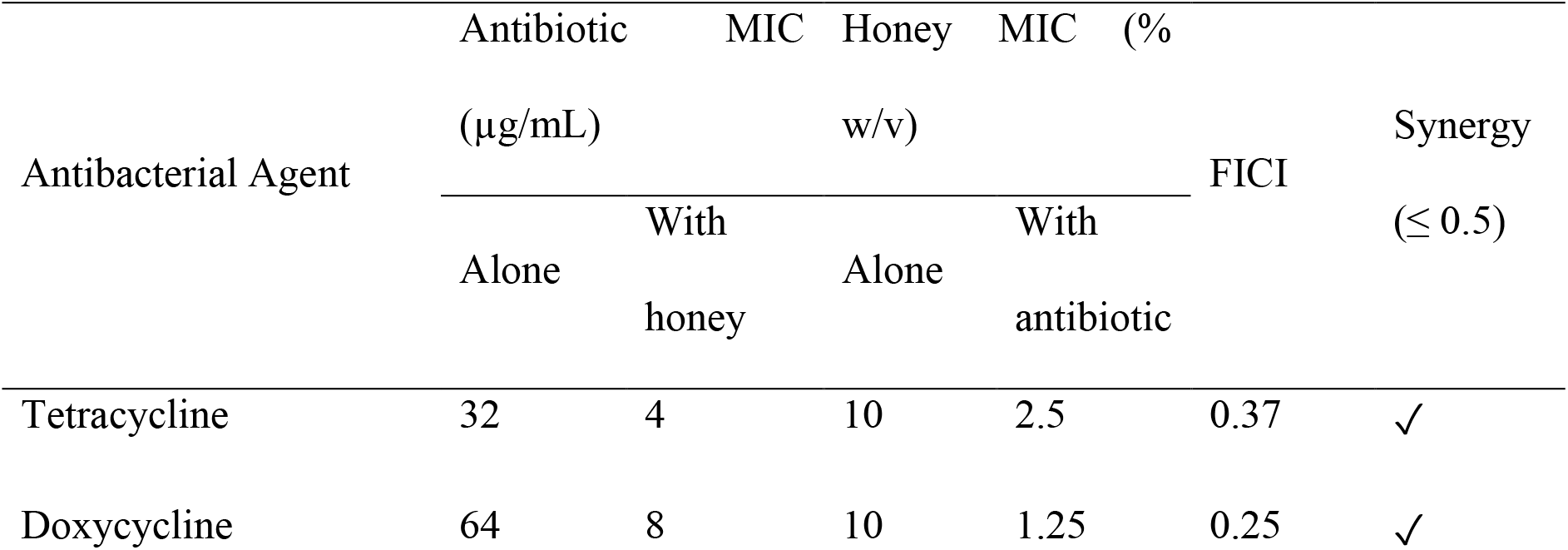
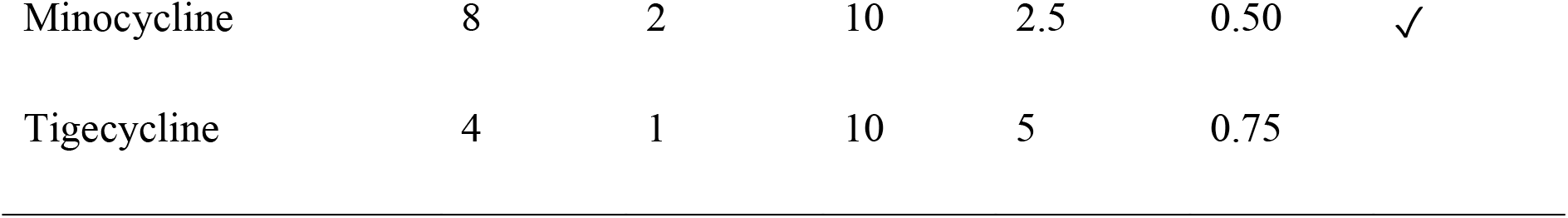
Summary of results from chequerboard analysis of combined effects of honey and tetracycline on P. aeruginosa PA14 growth.

We first explored the transcriptomic changes in *P. aeruginosa* induced by manuka honey. Manuka honey markedly affects the transcriptomic profile of *P. aeruginosa* compared to the untreated control (Figure 1A), with changes to the expression of 3177 of 5892 coding sequences (54%; false discovery rate threshold = 0.05). A similar number of genes were upregulated (n= 1646, representing 28% of all coding genes) versus downregulated (n=1531, or 26%). Analysis of only the genes with a log_2_FC of ≥ ±2 showed that 235 were differentially expressed, equivalent to 4% of all coding sequences. When this thresholding was applied, more genes were upregulated than downregulated (Figure 2A). Principle component analysis (PCA) confirmed that the effect of manuka honey on *P. aeruginosa* differed markedly relative to the untreated control (Figure 1B).

**Figure 1.**
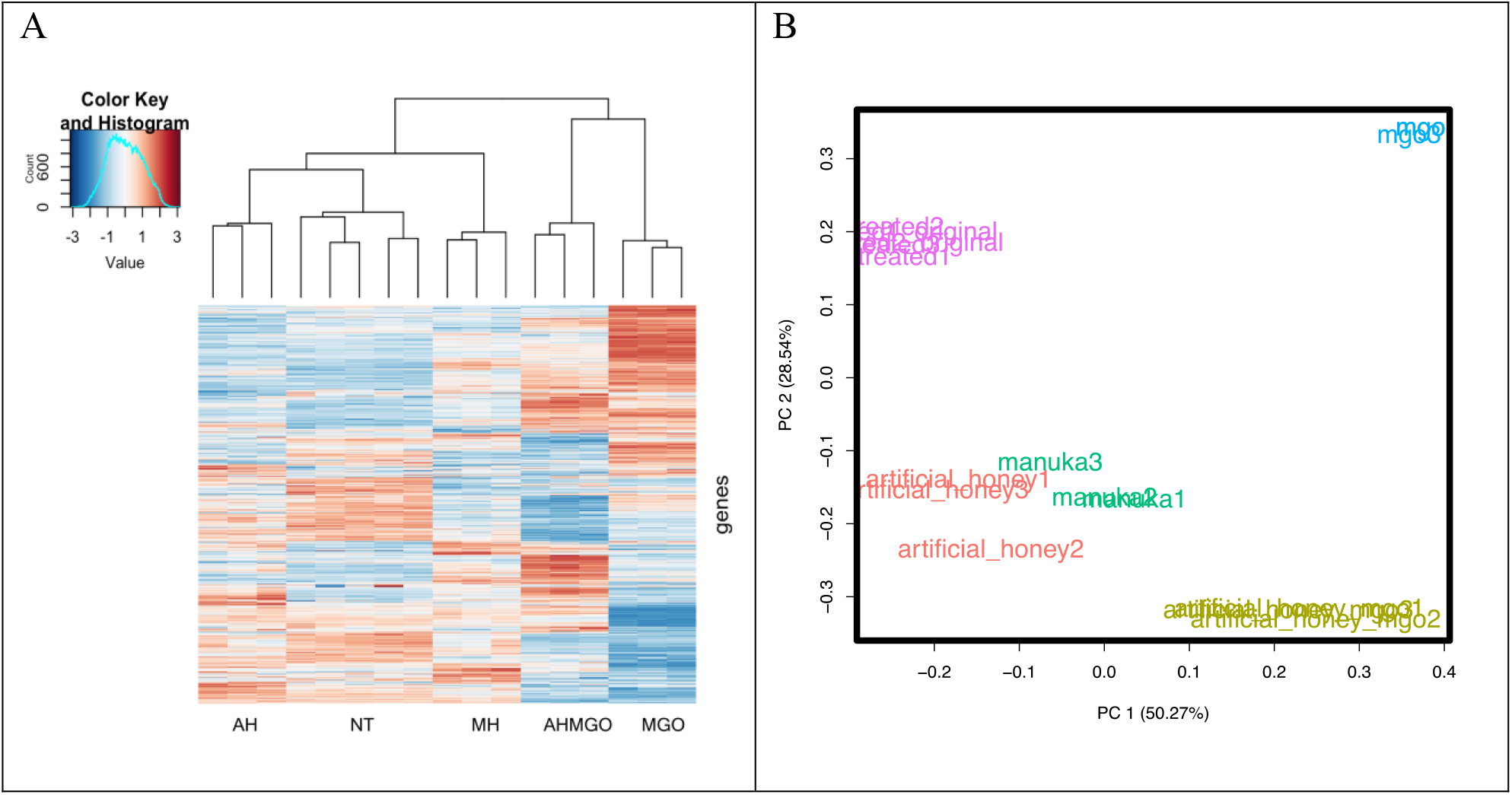
Transcriptional response of *P. aeruginosa* PA14 treated at mid-exponential phase with manuka honey and honey analogues for 30 minutes at
 0.5× MIC. (A) Clustered heatmap of relative expression of 5892 coding genes in *P. aeruginosa* across all treatments and a no-treatment control (NT). Treatments include whole manuka honey and its constituents: artificial honey (AH), artificial honey doped with methylglyoxal (AHMGO), and methylglyoxal (MGO) alone. RNA-seq was performed using three biological replicates for each treatment, and five for the no-treatment control. The clustered heatmap shows the Row Z-score (amount by which counts for a gene deviates in a specific sample from that gene’s average across all samples) and is clustered based on Euclidean measure and complete agglomeration (B) Bi-plot of the principal component analysis of normalised read counts for all treatments (manuka honey: green; MGO: blue; artificial honey: red; artificial honey + MGO: khaki) and the untreated control (purple), split into biological replicates.

**Figure 2.**
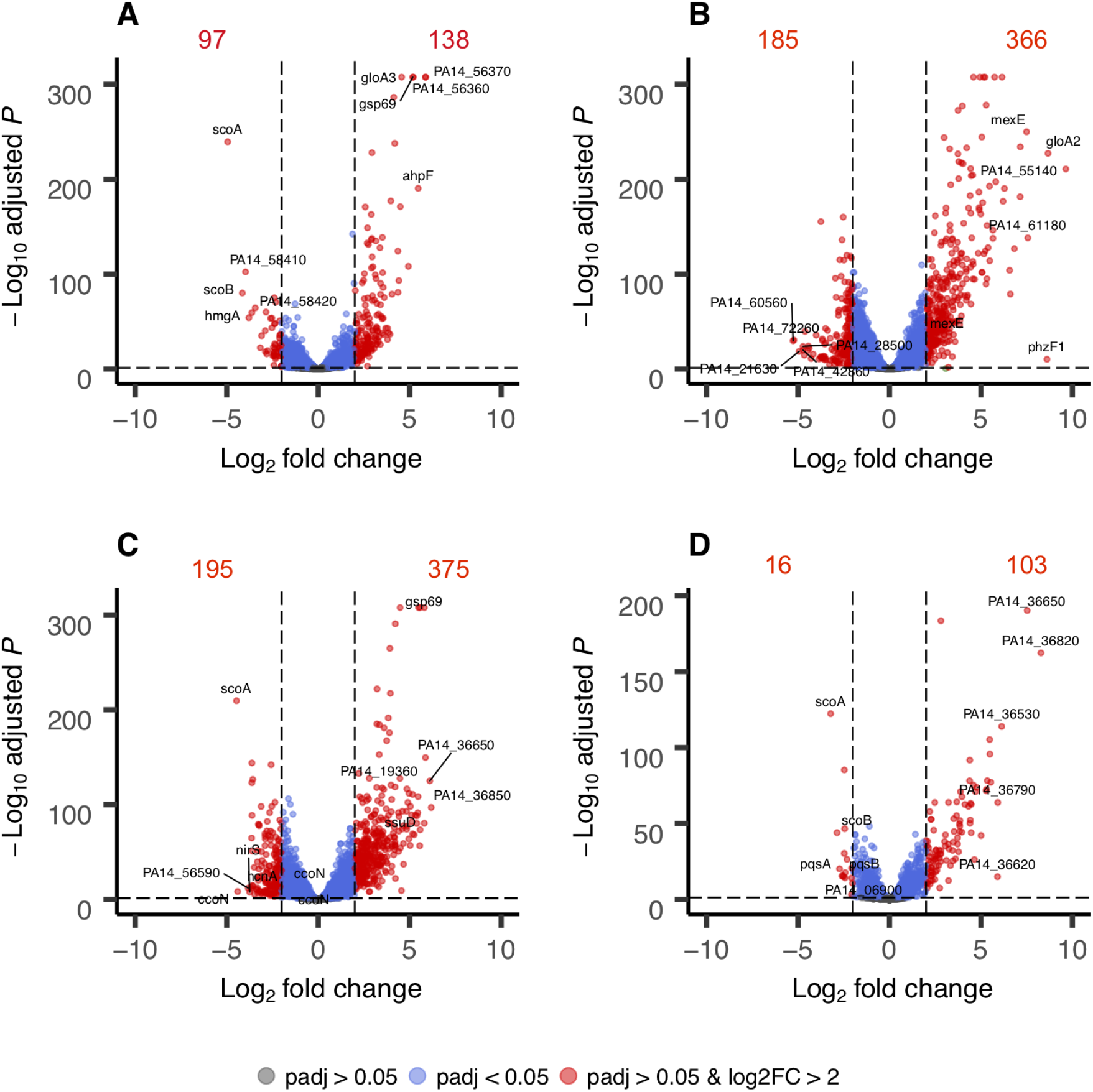
Summary of genome-wide expression changes in *P. aeruginosa* PA14 after treatment with A) manuka honey; B) MGO; C) AHMGO; and D) AH. The top 10 most differentially expressed genes are labelled in each plot. Grey dots indicate genes with no significant difference compared to the untreated control (p.adj > 0.05), blue dots indicate genes with a significant difference compared to untreated control (p.adj < 0.05) and red dots indicate genes with both a significant difference (p.adj <0.05) and log_2_FC > 2 compared to untreated control with numerical annotations to indicate the number of differentially expressed genes.

Genome-wide expression changes were visualised as volcano plots (Figure 2) to identify specific genes with large fold changes and statistical significance. Genes that were significantly differentially expressed (p.adj < 0.05) and above log_2_FC > 2 are presented in red and the five most up- and downregulated genes are labelled in each plot. Manuka honey treatment caused significant upregulation of 138 genes and downregulation of 97 genes (Figure 2A). In the manuka treated sample, the two genes in the PA14_56360-56370 operon were amongst the top five most upregulated (log_2_FC = 5.87 and 5.85, respectively). These genes encode hypothetical proteins that share homology with proteases from the DJ-1/PfpI family in *P. aeruginosa* PAO1, namely the oxidative stress response gene *ahpF* (log_2_FC = 5.46), the glyoxalase enzyme *gloA3* (log_2_FC = 5.19) and the aldo-keto reductase *gsp69* (log_2_FC = 5.16) (Figure 2A). The genes that had the largest downregulation following manuka treatment were the *scoAB* operon encoding CoA transferase subunits A and B (log_2_FC = − 4.94 and − 4.15 respectively), *hmgA* (log_2_FC = − 3.97) encoding a homogentisate 1-2-dioxygenase and a pair of genes in the PA14_58410-58490 operon encoding the outer membrane porin *opdP* (PA14_58410) (log_2_FC = − 3.79) and periplasmic ABC transporter *dppA4* (PA14_58420) (log_2_FC = − 3.62) (Figure 2A).

Differential expression of genes related to certain biological functions defined by PseudoCAP ^52^ were also analysed (Figure 3). PseudoCAP categories group genes based on experimental evidence of their involvement in a particular cellular function or their assignment to KEGG pathways participating in that function. The percentage of genes in each PseudoCAP classification that are differentially expressed (log_2_FC ≥±2 and p.adj ≤ 0.05) were used as an indication of the extent to which manuka honey affected particular biological functions in *P. aeruginosa*.

**Figure 3.**
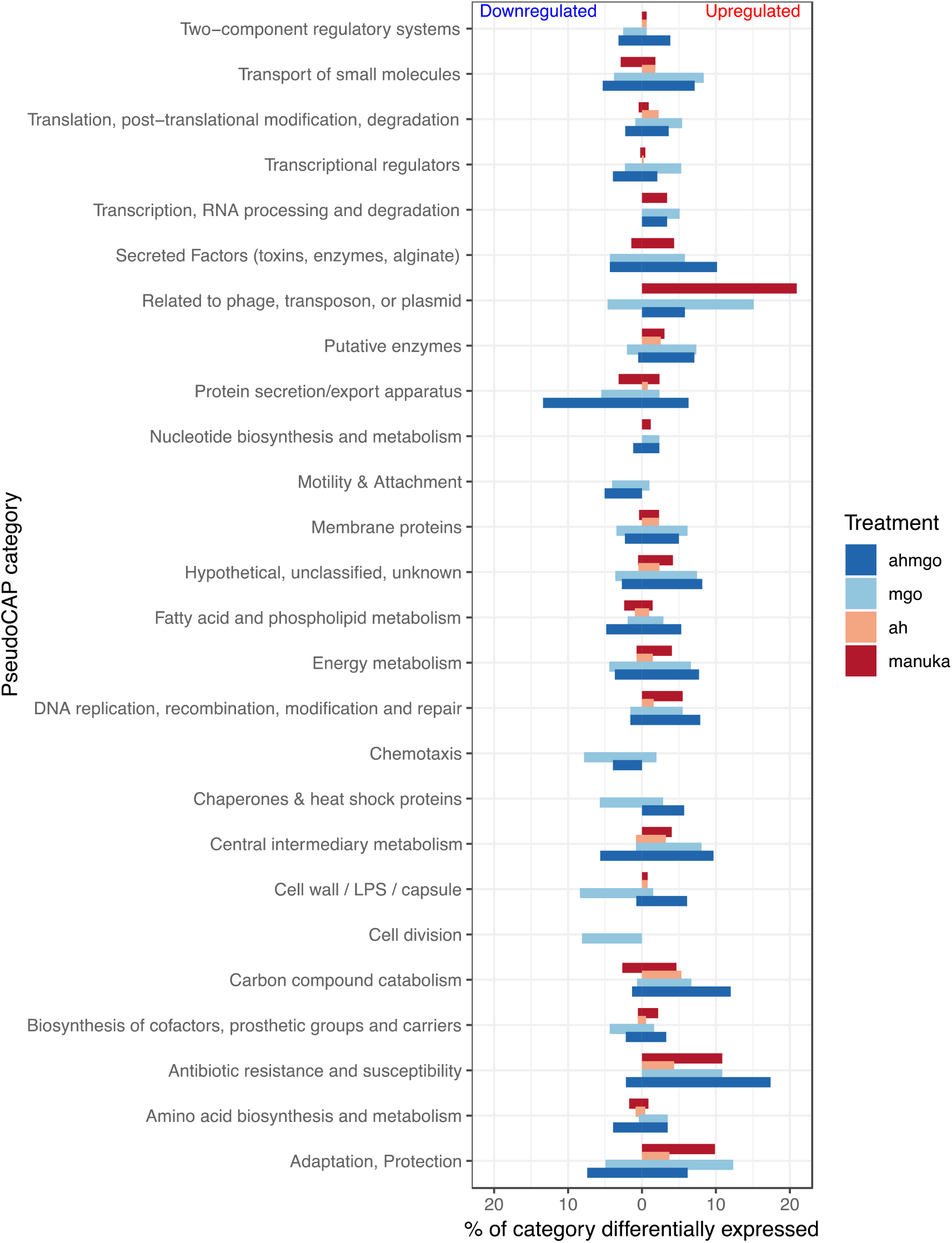
Percentage of genes of each PseudoCAP category which were differentially expressed (log_2_FC ≥±2 and p.adj ≤ 0.05) after treatment with manuka honey (dark-red), MGO (light-blue), AHMGO (dark-blue) and AH (light-red).

Overall, manuka honey had a major effect (upregulation) on the following categories: related to phage, transposon or plasmid; antibiotic resistance and susceptibility; and adaption and protection (Figure 3). Other categories upregulated by manuka honey in a sizeable but smaller manner were those relating to: transport of small molecules; transcription, RNA processing and degradation; secreted factors; putative enzymes; nucleotide biosynthesis metabolism; membrane proteins; energy metabolism; DNA replication, recombination, modification and repair; central intermediary metabolism; cell wall and biosynthesis of cofactors. In comparison, only a few processes were downregulated by manuka honey and these were: protein secretion; fatty acid and phospholipid metabolism, and amino acid biosynthesis and metabolism (Figure 3).

Functional groups corresponding to biological processes were manually curated and visualised as heatmaps (Figure 4). Functional groups were selected where several genes involved in a particular pathway or process were affected, and where at least two of those genes were amongst the 25 most differentially up- or downregulated. Manuka honey treatment induced the differential expression of genes involved in (but not limited to) quorum sensing (Figure 4A), the oxidative stress response (Figure 4B) and the SOS response (Figure 4C) and tailocin (sometimes referred to as pyocin) genes (Figure 4D; this is discussed later). To our knowledge, this is the first report of SOS induction by manuka honey in any microorganism. Our data suggesting that manuka honey affects quorum sensing via the downregulation of the *pqsABCDE* operon supports previous studies in *P. aeruginosa* PAO1 ^37^. Complementary techniques such as microarray analysis, genetic screens and proteomic approaches ^13, 29, 56, 57^ have shown honey can affect the expression of genes involved in the oxidative stress responses in *S. aureus* and *E. coli* and our findings indicate that this also occurs in *P. aeruginosa*.

**Figure 4.**
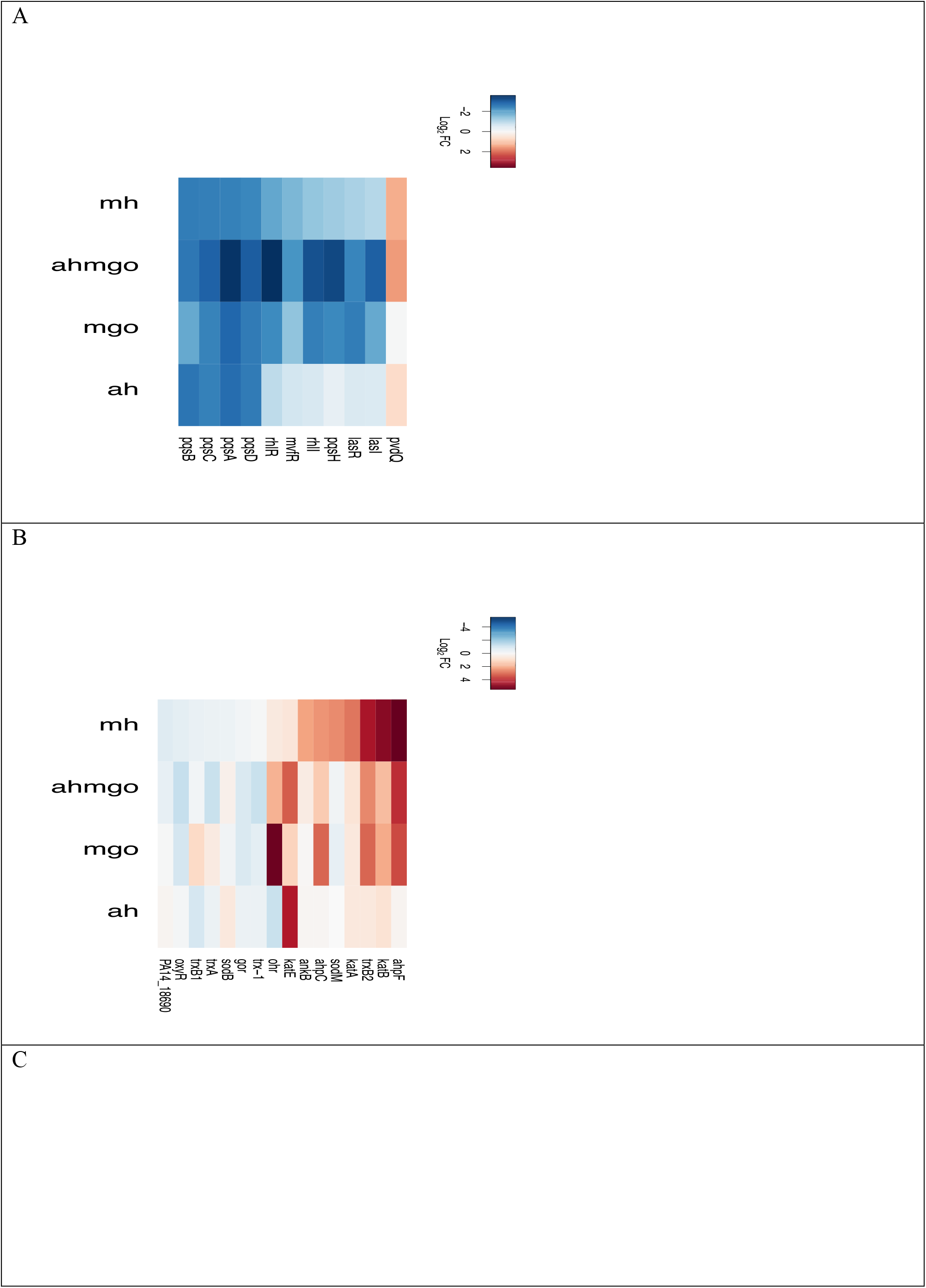
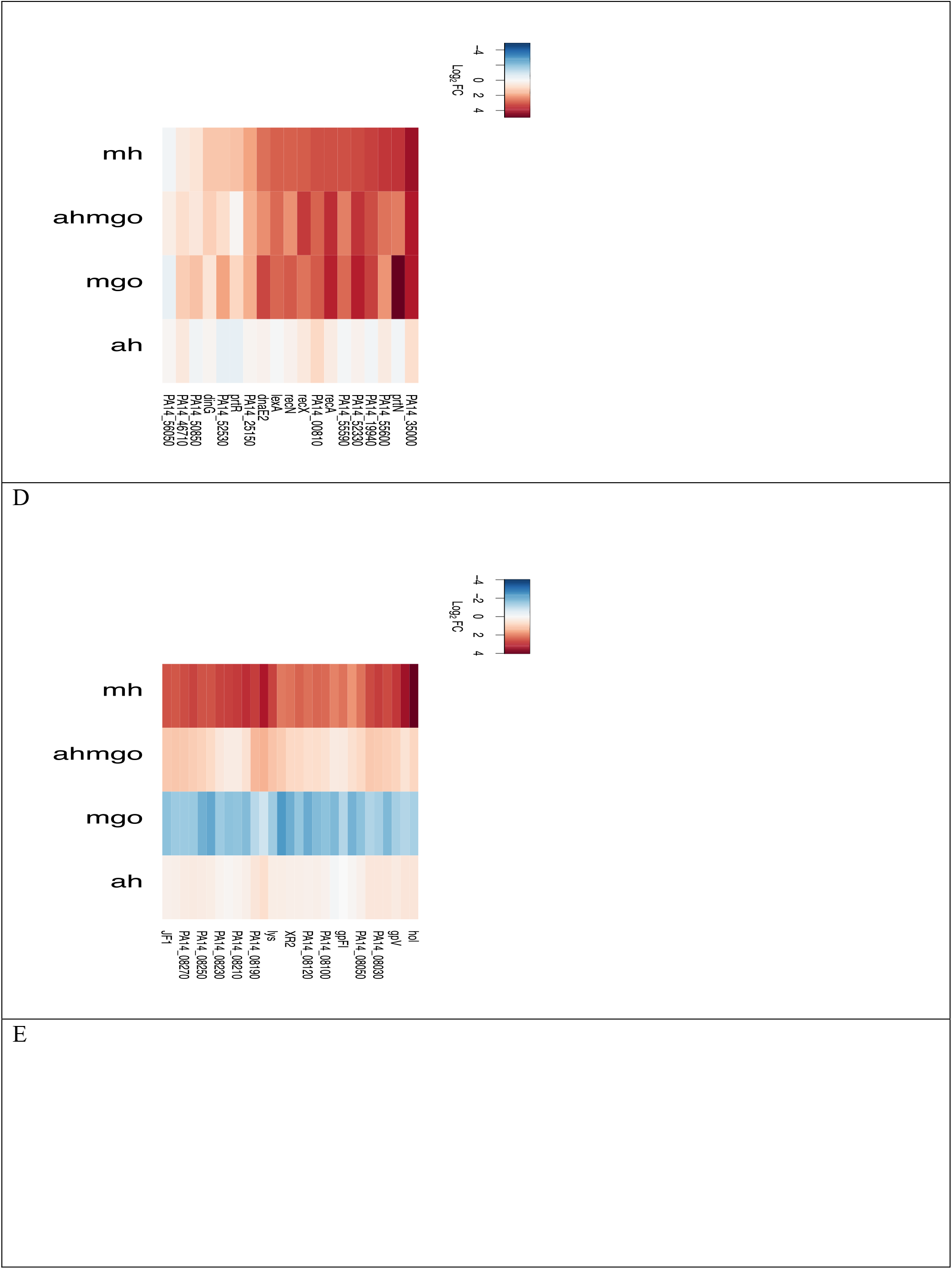
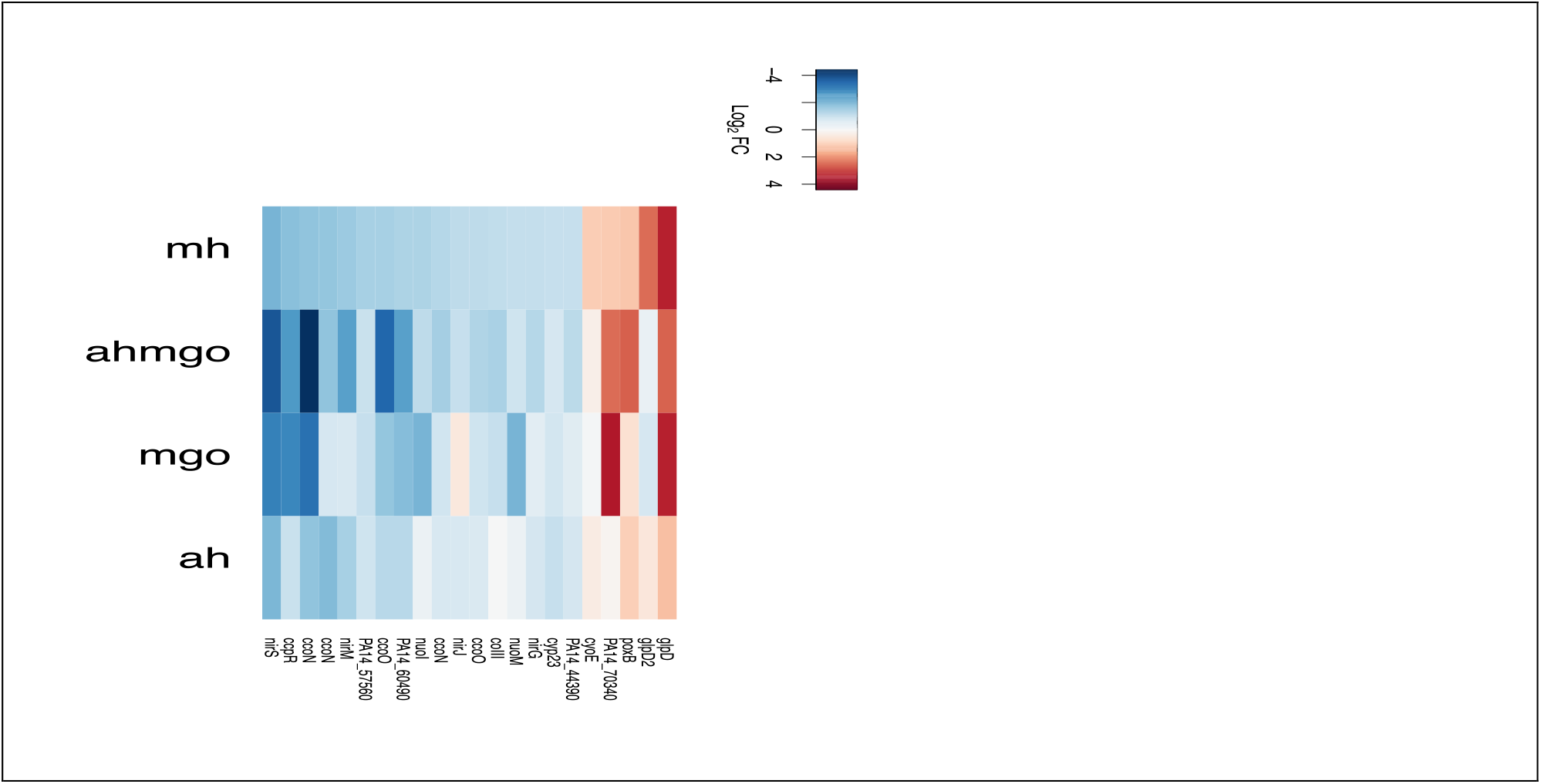
Heatmaps show log_2_FC data for *P. aeruginosa* PA14 treated at mid-exponential phase with manuka honey and honey analogues for 30 minutes at 0.5× MIC. A) quorum sensing genes B) oxidative stress response genes C) SOS response genes, D) tailocin genes, E) respiration genes for each treatment.

In order to explore whether oxidative stress responses were due to the generation of reactive oxygen species (ROS), thought to be a common killing factor of many antimicrobials, we performed MIC assays under anaerobic conditions where ROS formation is impeded ^58–60^. There was no difference in the MIC of manuka honey under aerobic versus anaerobic conditions, suggesting that ROS (and related oxidative stress) is not the only contributor to the antimicrobial mechanism of action. We have also previously demonstrated that exponentially growing *P. aeruginosa* PAO1 cells had condensed chromosomes after treatment with 4% w/v manuka honey, suggesting that it inhibits DNA replication in these cells ^22^. DNA degradation by oxidative damage would result in dispersed chromosomes rather than condensed ones, indicating that oxidative stress is not the mechanism of death in manuka honey treated *P. aeruginosa*.

### Can the transcriptomic effects of manuka honey on *P. aeruginosa* be accounted for solely by its key components?

The transcriptomic effects of manuka honey on *P. aeruginosa* appear to be greater than the sum of its parts, MGO and sugar, although there were many similar changes observed. Treatment with manuka honey induced transcriptional changes resulting in a unique gene expression profile when compared to the profiles of *P. aeruginosa* treated with the major components individually (Figure 1). Hierarchical clustering analysis of RNA-Seq data revealed that the manuka honey (MH) gene expression profile was most similar to AH and most different to MGO alone. The combination of AHMGO was more similar to MH than MGO alone (Figure 1A) and this is supported by PCA (Figure 1B).

Analysis of individual gene changes showed that while many genes are significantly differentially expressed (log_2_FC > 2; p.adj < 0.05) across all treatments, MGO treatment resulted in the highest number of differentially expressed genes overall (Figure 2). This could be, in part, due to the higher concentration of MGO; because using 0.5× MIC across all treatments meant that the inhibitory effect on cells was the same but the final MGO concentration was different under each condition. There are several genes amongst the five most up- or downregulated ones common across multiple treatments (MH, MGO, AHMGO) such as *ahpF*, which has been previously identified and is known to play an important role in the response to oxidative species (Figure 2). The genes *scoA* and *gsp69* were in the ten most significantly differentially expressed genes across multiple treatments (*scoA* in MH, AHMGO and AH, *gsp69* in MH and AHMGO). The gene *gsp69* encodes a probable oxidoreductase with homology to aldo-keto reductases (AKRs) in *E. coli* ^61^. AKRs are capable of detoxifying MGO by reducing methylglyoxal to hydroxyacetone (the monomeric form of DHA found in manuka honey) using NADPH as a co-factor ^62–65^. Only six genes were differentially expressed across all treatments (Figure 5). These included the PQS quorum sensing genes *pqsACDE* and the gene immediately downstream from this operon, *phnA*. All treatments affect the PQS quorum sensing system (Figure 4A) and while this is consistent with previous reports ^37, 39^, it has not been reported for treatment with MGO alone.

**Figure 5.**
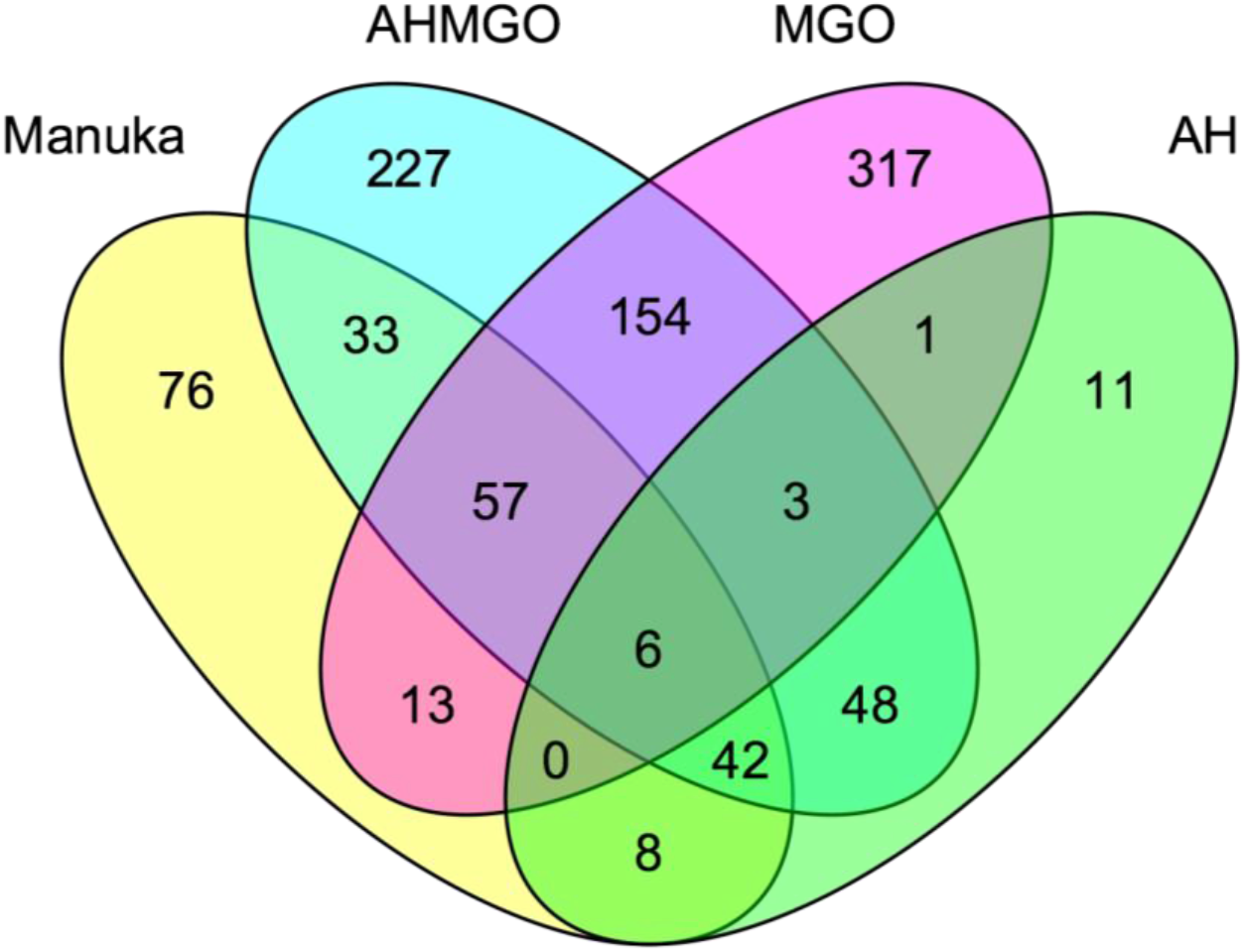
Venn diagram of differentially expressed genes (log_2_FC ≥ 2, p.adj ≤ 0.05) in P. aeruginosa PA14 after treatment with (yellow) manuka honey (yellow), AHMGO (blue), MGO (pink) and AH (green) at 0.5× MIC.

In general, manuka honey affected the same processes as AHMGO and MGO, however not always to the same extent, nor in the same direction (Figure 3). Many of the PseudoCAP categories shown as being affected by honey, including secreted factors, protein secretion, energy metabolism, and DNA replication and adaption, were similarly affected by MGO and AHMGO treatment, but not AH (Figure 3). This suggests that MGO contributed, at least in part, to these gene expression changes. The data also indicate that AH actually has little contribution to the action of manuka honey in terms of the gene expression changes in the biological process and pathways of *P. aeruginosa*, but the sugar component may be necessary to facilitate MGO activity (Figure 3 and Figure 4).

There were certain categories that were differently affected by manuka honey relative to its major components, for example genes in the ‘transport of small molecules’ category were mainly downregulated by manuka honey whereas AHMGO and MGO seemed to induce both up- and downregulation. The significantly higher number of differentially expressed genes in AHMGO and MGO treatment may be a downstream effect of the higher degree of differential expression of genes in the ‘transcriptional regulators’ category (Figure 3), which includes transcriptional regulators such as *lasR, rhlR, algQ*, and *mvfR* (also known as *pqsR*). All of these genes are global regulators controlling transcription of large sets of genes across the *P. aeruginosa* genome in response to different stimuli ^66^.

Like manuka, the expression of genes in the oxidative stress response was also affected by AHMGO and MGO. This data is congruent with the expression data of genes involved in the SOS response, where AHMGO and MGO induced strong upregulation in a wide range of genes involved in SOS, notably *recA* and *lexA* (Figure 4). MGO is known to cause damage to DNA by modification of guanine bases ^27, 67, 68^, and has been reported to inhibit the initiation of DNA replication, causing double stranded breaks in DNA that induces DNA repair ^26^. MGO treatment has been shown to induce the SOS response in *B. subtilis* ^69^, therefore the strong upregulation of genes in the SOS response by both MGO and AHMGO is not surprising.

Curiously, our data shows that manuka honey induces comparable expression of SOS response genes to MGO and AHMGO treated cells despite containing a much lower concentration of MGO. Previous research shows that the expression of SOS genes reduces over time after initial exposure to MGO and this is thought to be due to the initial transient depletion of glutathione (GSH) which is required for the function of the GSH-dependent glyoxalase systems of *gloA* genes ^26^. While it is clear that MGO plays a role here, the upregulation of SOS genes by manuka honey cannot be solely attributed to this component.

Whilst similarities across the treatments were seen, we identified 76 genes as being uniquely differentially expressed by manuka honey (Figure 5). These genes included phage-related genes (Figure 3), such as the chromosomally encoded tailocin genes *hol* and *lys* (Figure 4D), which are involved in explosive cell lysis, mediated though the tailocin pathway and dependent on endolysin (*lys*). We also saw significant gene expression changes in haem oxygenase *nemO*, oxidative stress response genes *sodM*, *ankB* and *katA* and metabolic genes *fumC1* and *glpD2*. Uniquely downregulated genes include those encoding ABC transporters, *ybeJ*, *gltJK* and *dppD*, metabolic genes *atoB*, *braE*, *maiA*, *fahA* and *gnyB* and the cytoplasmic potassium transporter K^+^ binding and translocating subunit *kdpA*.

We tested the susceptibility of single-gene knock-out mutants (n= 23) of *P. aeruginosa* that were either highly or uniquely differentially expressed after manuka treatment (Table S3). One mutant, Δ*gloA3* (encoding a glyoxalase enzyme for MGO detoxification), showed increased susceptibility (MIC = 5%) relative to the wild-type (MIC = 10%) (Table S3), suggesting that this gene is important for the survival of *P. aeruginosa* in the presence of honey. The Δ*gloA3* strain is deficient in lactoylglutathione lyase, one of the three redundant enzymes required for the conversion of MGO and glutathione to lactoylglutatione suggesting that lactoylglutathione lyase plays a role in the antimicrobial action of manuka. However, it is still unclear whether this is solely due to its capacity to detoxify MGO or other downstream effects. The chemical complexity of honey would suggest that it targets multiple pathways or proteins therefore a single mutation may not lead to a change in MIC. This is consistent with the inability of bacteria to develop resistance to honey ^7^.

One of the most perturbed pathways for manuka honey treated cells related to aerobic respiration, for example, *nemO*, *phuT* and *phuS* (Figure 5). PhuST can maintain iron homeostasis by binding haem and either stores it or transferring it to NemO which is then able to liberate iron ^70^. Haem is a cofactor of cytochromes and acts as the electron shuttle for many enzymes in the electron transport chain, playing a critical role in cellular respiration ^71^. Combined with the expression levels of genes involved in the electron transport chain and central carbon metabolism (Figure 4E), along with the unique expression of the cytoplasmic membrane depolarising gene *hol* ^72, 73^, we hypothesised that manuka honey may be affecting the proton motive force (PMF) of *P. aeruginosa*.

### Collapse of the proton motive force: a unique contributor to the antimicrobial activity of manuka honey?

We applied two independent approaches to investigate the impact of manuka on the PMF. To examine directly whether compounds in manuka facilitate the passage of protons across biological membranes, we used liposomes loaded with the pH sensitive dye pyranine allowing detection of proton movement across the liposome lumen using fluorescence. Liposomes were formed in buffer containing only potassium salts at pH 7.0 and then diluted into buffer containing only sodium at pH 7.0. The addition of a low concentration of the potassium ionophore valinomycin, allowed potassium to move down its concentration gradient out of the liposomes, generating an outside positive electrical gradient. Subsequent addition of manuka caused a rapid drop in the internal pH indicating proton movement into the liposomes (Figure 6A). The proton movement was dependent on the establishment of an electrical gradient, since no pH change was observed in the absence of valimomycin. No significant change in pH was observed in experiments using AH, AHMGO or MGO alone, suggesting that a unique component manuka honey is required. This component appears to be acting as a novel protonophore that facilitated the passage of protons down an electrical gradient across a biological membrane (Figure 6A).

**Figure 6.**
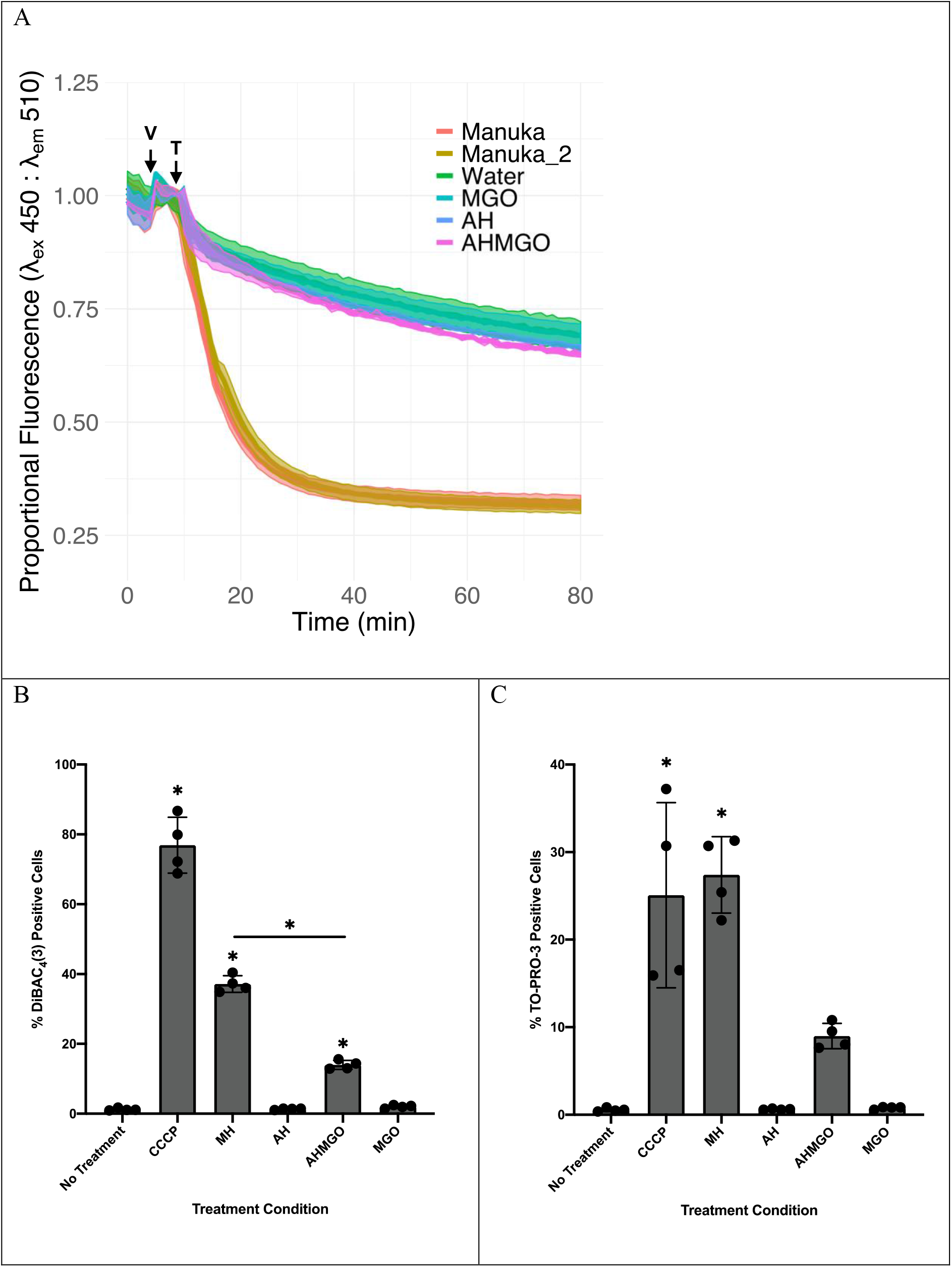
The effect of manuka honey on membrane potential measured as A) approximate internal pH of liposomes loaded with pH sensitive pyranine dye after treatment with 1% w/v artificial honey (blue), 1% w/v MGO (teal), 1% w/v AHMGO (purple), 1% manuka honey (orange), 1% manuka honey (tan). Fluorescence intensity of pyranine at λ_ex_ 450 decreases with decreasing pH. A decrease in proportional fluorescence is indicative of a decrease in internal pH. ‘V’ indicates addition of valinomycin to establish electrical gradient and ‘T’ indicates time at which treatment was applied, B) flow cytometry quantification of the percentage of DiBAC_4_(3) positive exponential phase *P. aeruginosa* PAO1-EcPore cells treated for 2 hours with 100 µM CCCP, 10% w/v manuka honey, 10% AH, 10% w/v AHMGO, 10% MGO and no treatment control. The effect of manuka honey on membrane permeability measured as C) flow cytometry quantification of the percentage of TO-PRO^®^-3 positive exponential phase *P. aeruginosa* PAO1-EcPore cells treated for 2 hours with 100 µM CCCP, 10% w/v manuka honey, 10% AH, 10% w/v AHMGO, 10% MGO and no treatment control. A one-way ANOVA followed by Dunnett’s multiple comparison post hoc test was used to determine statistically significant differences between each treatment and the no treatment control (* = *P* < 0.05), a one-way ANOVA followed by Bonferroni’s multiple comparison post hoc test was used to determine statistically significant differences between manuka and AHMGO treated cells (* = *P* < 0.05).

To validate whether manuka honey can induce membrane depolarisation in live *P. aeruginosa* cells, we used flow cytometry with DiBAC_4_(3), a fluorophore subject to selective uptake in depolarised cells (representative plots shown in Figure S3). We assessed the number of cells with depolarised membranes after treatment (2 hours) with manuka and its key components. CCCP (100 µM), a PMF uncoupler, was included as a positive control (Figure 6B). Measurement of membrane potential in *P. aeruginosa* is complicated by outer membrane exclusion of fluorophores and EDTA pre-treatment is often used to increase dye uptake in Gram-negatives ^74, 75^. However, this induced wide-spread membrane depolarisation making negative and positive controls indistinguishable (data not shown). A well characterised ‘hyperporinated’ *P. aeruginosa* PAO1 strain expressing a chromosomally encoded gene for a modified *E. coli* siderophore uptake channel ^76^ was used to overcome these limitations.

Manuka honey induced significant membrane depolarisation in *P. aeruginosa*, unlike MGO and AH (Figure 6B). Whilst AHMGO induced significant membrane depolarisation relative to the untreated control, this was at levels significantly lower than manuka honey. Exchange of protons across lipid bilayers by manuka honey (Figure 6A) suggests a dissipation of the PMF and is consistent with our data showing an overall collapse of PMF in *P. aeruginosa* after treatment with manuka (Figure 6B). However, manuka treatment resulted in an increased number of cells positive for TO-PRO-3 fluorescence, indicating membrane permeabilisation. This strongly suggests that the depolarisation observed in *P. aeruginosa* cells is a result of damage to the cytoplasmic membrane (Figure 6C).

Because of the observation of membrane permeabilisation and depolarisation, we hypothesised that manuka honey may affect the activity of antibiotics, for example tetracyclines, to which *P. aeruginosa* is innately resistant. Tetracyclines inhibit the binding of aminoacyl-tRNA to the mRNA translation complex ^77^ and a major mechanism of tetracycline resistance is through cytoplasmic membrane drug transporters, which require PMF for drug exportation ^77, 78^. We expect manuka honey treatment could increase tetracycline uptake due to increased permeabilisation and reduce efflux as a result of PMF collapse and thus enhance the potency of these antibiotics.

Accordingly, four tetracyclines were chosen for assessment of synergistic interaction with manuka honey by chequerboard assays, among which tetracycline, doxycycline and minocycline are known substrates to Tet efflux pumps (TetA/B) ^77^ and some resistance-nodulation-division (RND) family transporters (MexAB-OprM, MexXY-OprM and MexEF-OprN) ^79, 80^, but tigecycline is not recognised by Tet transporters and has a much weaker interaction as a substrate to the RND pumps ^81^. The functionality of the RND pumps is also membrane potential dependent. Consistent with our hypothesis, apart from tigecycline, manuka honey had strong synergy with the tetracycline antibiotics (Table 2). Furthermore, the synergy is positively correlated with the MICs of the tetracyclines (Table 2), suggesting that manuka honey is effective to restore tetracycline antibiotic potency against the bacterial strains that would otherwise be resistant. Tetracyclines have other resistance determinants, such as ribosomal protection proteins and enzymatic inactivation ^77^. We cannot exclude the possibility that the cause of the tetracycline-manuka synergy is more than membrane depolarisation and permeabilization, but this is beyond the scope of current study.

The PMF is an attractive target for antimicrobial therapy as it is a fundamental process in energy generation for bacteria. A collapse in the PMF impedes the ability of bacteria to generate energy required to drive processes necessary for resistance to antibiotics, for example, detoxification of tetracycline by PMF-driven multidrug resistant efflux pumps^78^. Our data suggests that manuka honey is collapsing the PMF in *P. aeruginosa* (Figure 6), and that this may be a biophysically driven mechanism such as damage to the cytoplasmic membrane, and is independent of proteins involved in the electron transport chain. Previous reports have shown that manuka honey acts synergistically with tetracycline against *S. aureus* ^82^ but only additively against *P. aeruginosa* ^83^, however, our data suggest manuka honey also acts synergistically with tetracycline against *P. aeruginosa* (Table 2). *P. aeruginosa* is intrinsically resistant to tetracycline due to drug efflux mediated through PMF dependant MexAB-OprM and MexXY-OprM RND multidrug efflux pumps ^84^. Collapse of the PMF would also impair the function of proton dependent efflux systems, which may explain the synergistic interaction of manuka honey and tetracyclines. This data suggests a role for manuka honey as a therapeutic adjuvant potentially restoring the therapeutic utility of antimicrobials no longer used to treat *P. aeruginosa* infections or open up new treatment options for topical *P. aeruginosa* infections.

## Conclusions

This study is the first to use a global transcriptomic approach, RNA-seq, in combination with classic microbiology techniques, to investigate the effects and antibacterial mechanism of action of manuka honey and its key antibacterial components against *P. aeruginosa*. We demonstrate that: (1) manuka honey induces wide-spread transcriptional changes and affects many biological processes; (2) these changes are not wholly explained by its key components, sugar and MGO, either alone or in combination; (3) MGO, widely accepted as the single most important antibacterial component of manuka honey, does not account for its total activity against *P. aeruginosa* (4) collapse of the proton motive force and membrane permeabilisation may be a key contributor to the unique antimicrobial activity of manuka honey.

## Acknowledgements

We thank Comvita New Zealand for the supply of the manuka honey sample. This work was supported by a UTS Doctoral Scholarship awarded to Daniel Bouzo. Flow cytometry was performed at the UTS Microbial Imaging Facility. We also thank Prof Merilyn Manley-Harris for advice on the formulation of artificial honey and Prof. Helen Zgurskaya for providing the hyperporinated *P. aeruginosa* strain.

## Author contributions

DB, EH, KH, NC contributed to the conception and design of the work. DB, NC, EH, LL, KH, GB, JL, AB contributed to the acquisition, analysis and interpretation of data for the work. The paper was written by DB and NC, and critically revised by EH, LL, KH, IP, AB, CW.

## Data availability Statement

All data generated and analysed during this study are included in this published article, its Supplementary Material files, and deposited in the Gene Expression Omnibus (GEO) data with the accession number GSE142448 (https://www.ncbi.nlm.nih.gov/geo/query/acc.cgi?acc=GSE142448).

## Supplementary Material (For Publication)

**Table S1.** Strains used in this study.

| Strain | Relevant characteristics | Source or reference |
| --- | --- | --- |
| *P. aeruginosa* |  |  |
| PAO1 | Wildtype | <sup>85</sup> |
| PA14 | Wildtype | <sup>86</sup> |
| PAO1Δ*recA* | *recA* deletion mutant of PAO1 | <sup>73</sup> |
| PAO1Δ*lys* | *lys* deletion mutant of PAO1 | <sup>73</sup> |
| PAO1Δ*prtN* | *prtN* deletion mutant of PAO1 | <sup>73</sup> |
| PAO1-LAC-EcPore | ‘hyperporinated’ PAO1, a chromosomally encoded modified *E. coli* FhuA siderophore uptake channel (EcPore) with an IPTG inducible promoter. | <sup>76</sup> |
| PAO1-LAC | Empty expression cassette | <sup>76</sup> |
| ΔPA14_03930 | PA14_03930::MAR2xT7 | <sup>87</sup> |
| ΔPA14_06510 | PA14_06510::MAR2xT7 | <sup>87</sup> |
| ΔPA14_18750 | PA14_18750::MAR2xT7 | <sup>87</sup> |
| ΔPA14_18760 | PA14_18760::MAR2xT7 | <sup>87</sup> |
| ΔPA14_18780 | PA14_18780::MAR2xT7 | <sup>87</sup> |
| ΔPA14_24940 | PA14_24940::MAR2xT7 | <sup>87</sup> |
| ΔPA14_32380 | PA14_32380::MAR2xT7 | <sup>87</sup> |
| ΔPA14_33520 | PA14_33520::MAR2xT7 | <sup>87</sup> |
| ΔPA14_33970 | PA14_33970::MAR2xT7 | <sup>87</sup> |
| ΔPA14_38040 | PA14_38040::MAR2xT7 | <sup>87</sup> |
| ΔPA14_38640 | PA14_38640::MAR2xT7 | <sup>87</sup> |
| ΔPA14_38660 | PA14_38660::MAR2xT7 | <sup>87</sup> |
| ΔPA14_51830 | PA14_51830::MAR2xT7 | <sup>87</sup> |
| ΔPA14_51430 | PA14_51430::MAR2xT7 | <sup>87</sup> |
| ΔPA14_51600 | PA14_51600::MAR2xT7 | <sup>87</sup> |
| ΔPA14_53290 | PA14_53290::MAR2xT7 | <sup>87</sup> |
| ΔPA14_58000 | PA14_58000::MAR2xT7 | <sup>87</sup> |
| ΔPA14_58030 | PA14_58030::MAR2xT7 | <sup>87</sup> |
| ΔPA14_58450 | PA14_58450::MAR2xT7 | <sup>87</sup> |
| ΔPA14_61020 | PA14_61020::MAR2xT7 | <sup>87</sup> |
| ΔPA14_62400 | PA14_62400::MAR2xT7 | <sup>87</sup> |
| ΔPA14_67500 | PA14_67500::MAR2xT7 | <sup>87</sup> |

**Table S2.** Concentrations (0.5 × MIC) of treatments applied to *P. aeruginosa* PA14 cells prior to RNA harvest and the respective MGO content at each concentration.

| Treatment | 0.5 × MIC (% w/v) | MGO content (ppm) |
| --- | --- | --- |
| Manuka honey | 5 | 45 |
| MGO | 27.5 | 247.5 |
| AHMGO | 10.75 | 96.75 |
| AH | 12.5 | 0 |

**Table S3.**
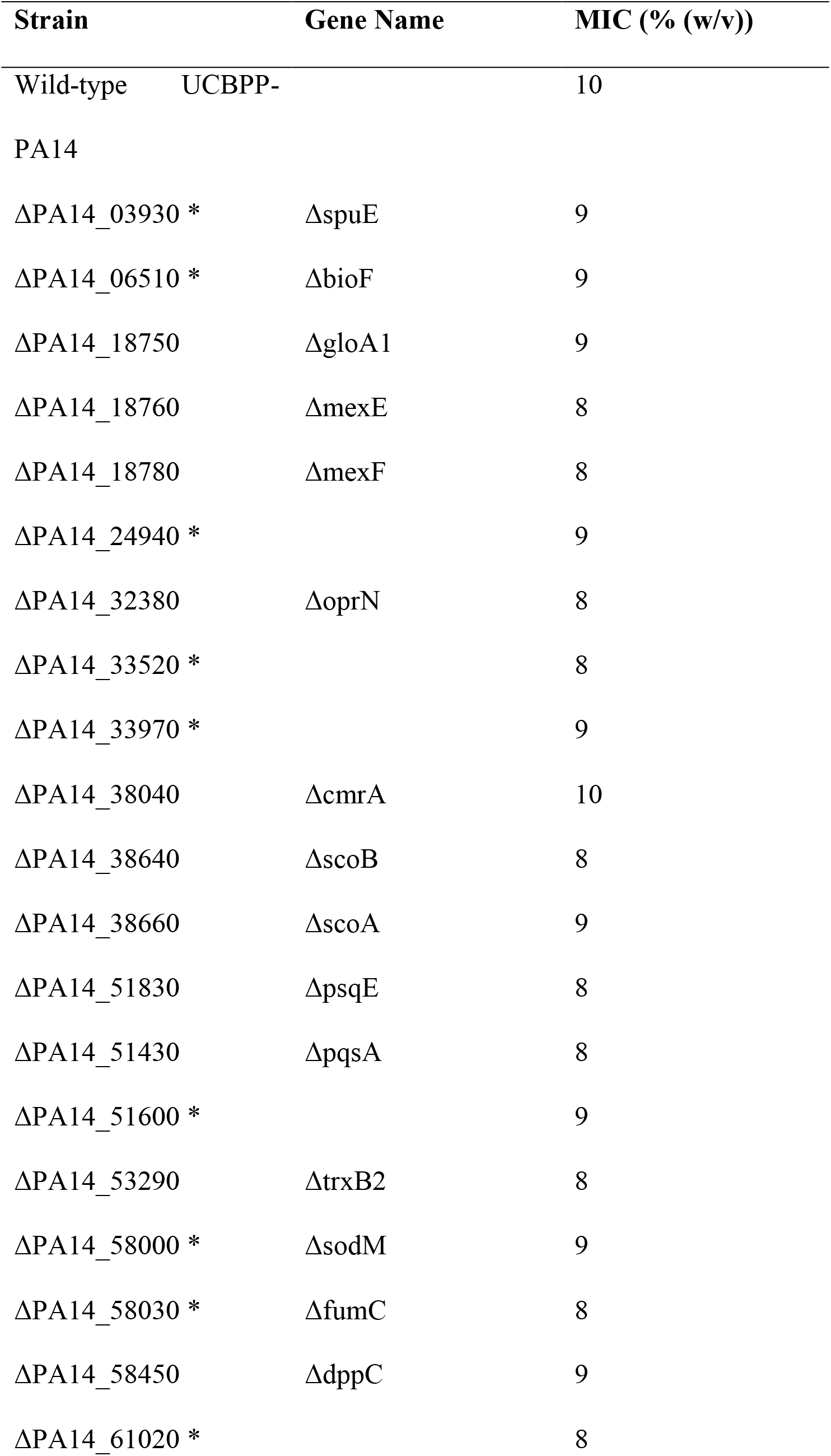
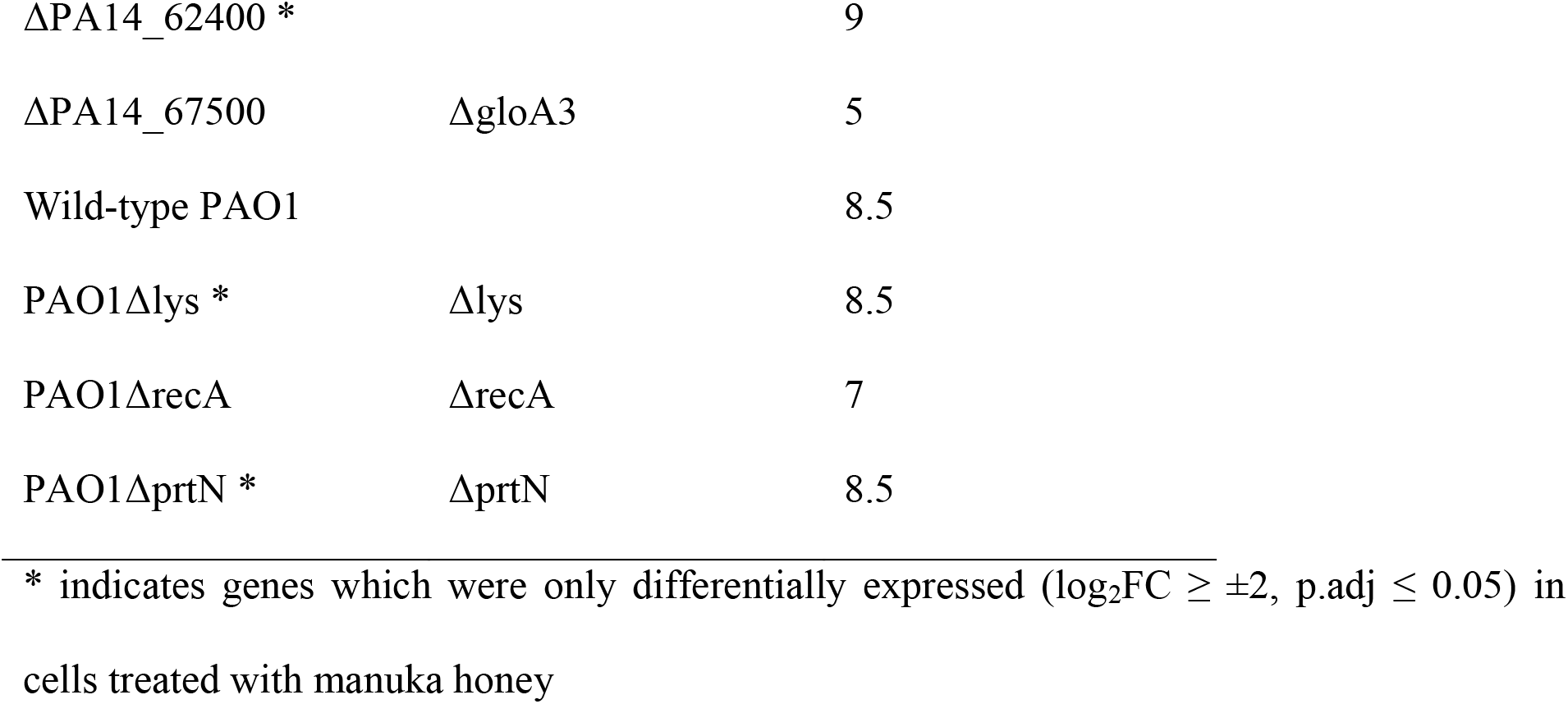
Susceptibility to manuka honey treatment for mutant strains.

**Table S4.**
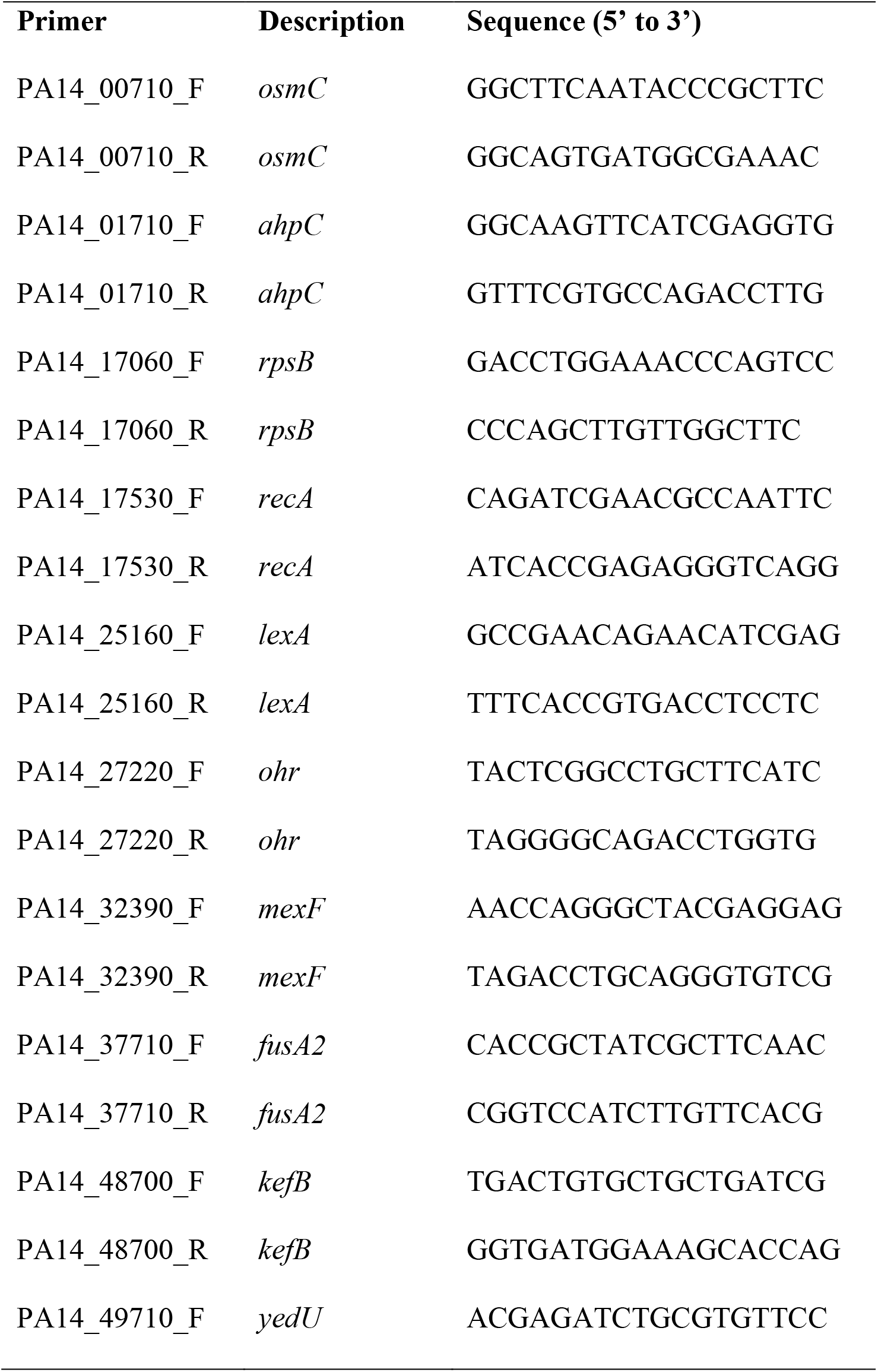
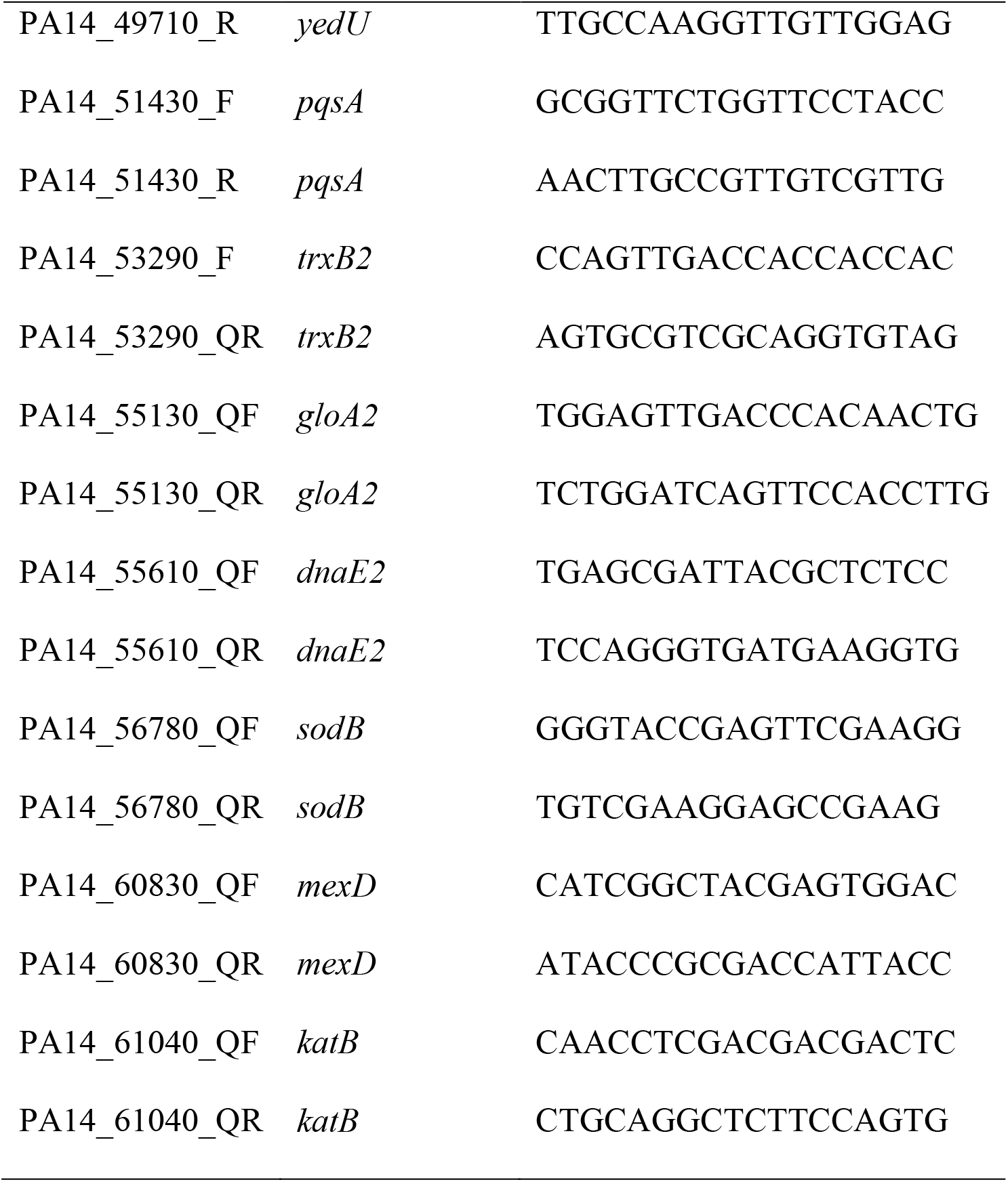
Primers used for RT-qPCR reactions.

**Fig. S1.**
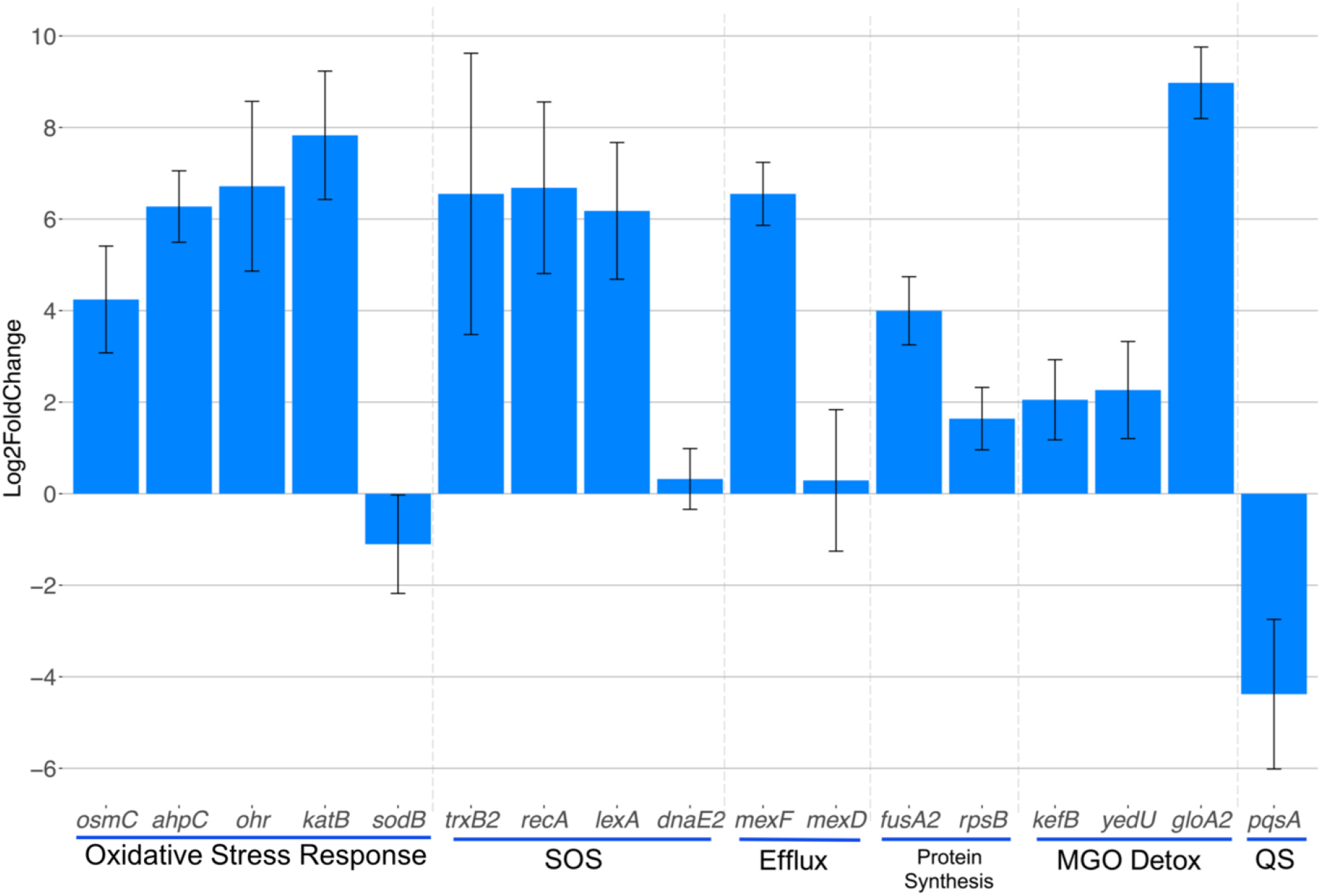
Changes in gene expression following treatment with 5% (0.5x MIC) manuka honey. Data are expressed as mean values ± SD from one technical replicate, conducted using three biological replicates.

**Fig. S2.**
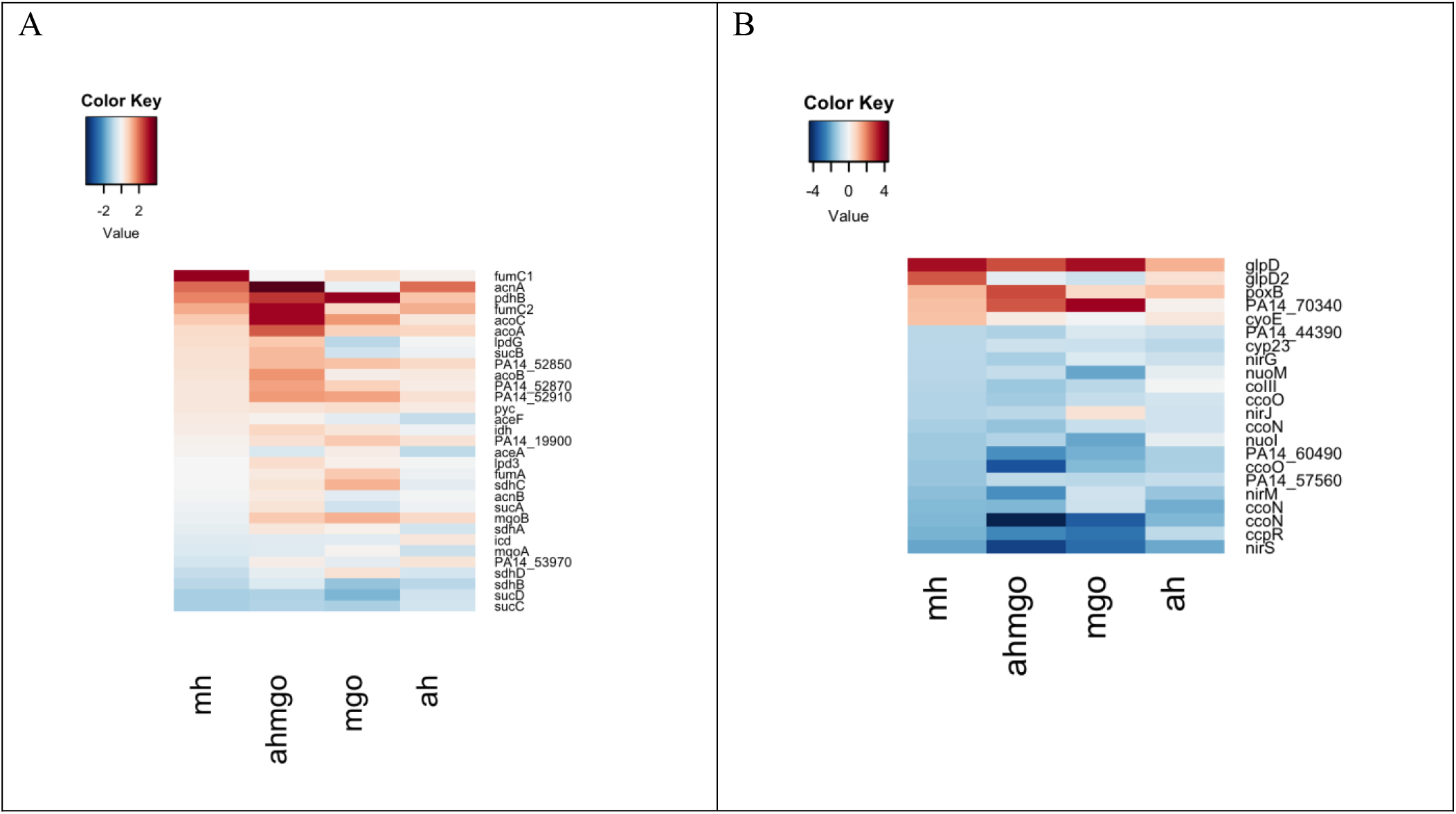
Heatmaps of log_2_FC of genes involved in A) central carbon metabolism and B) aerobic respiration

**Fig. S3.**
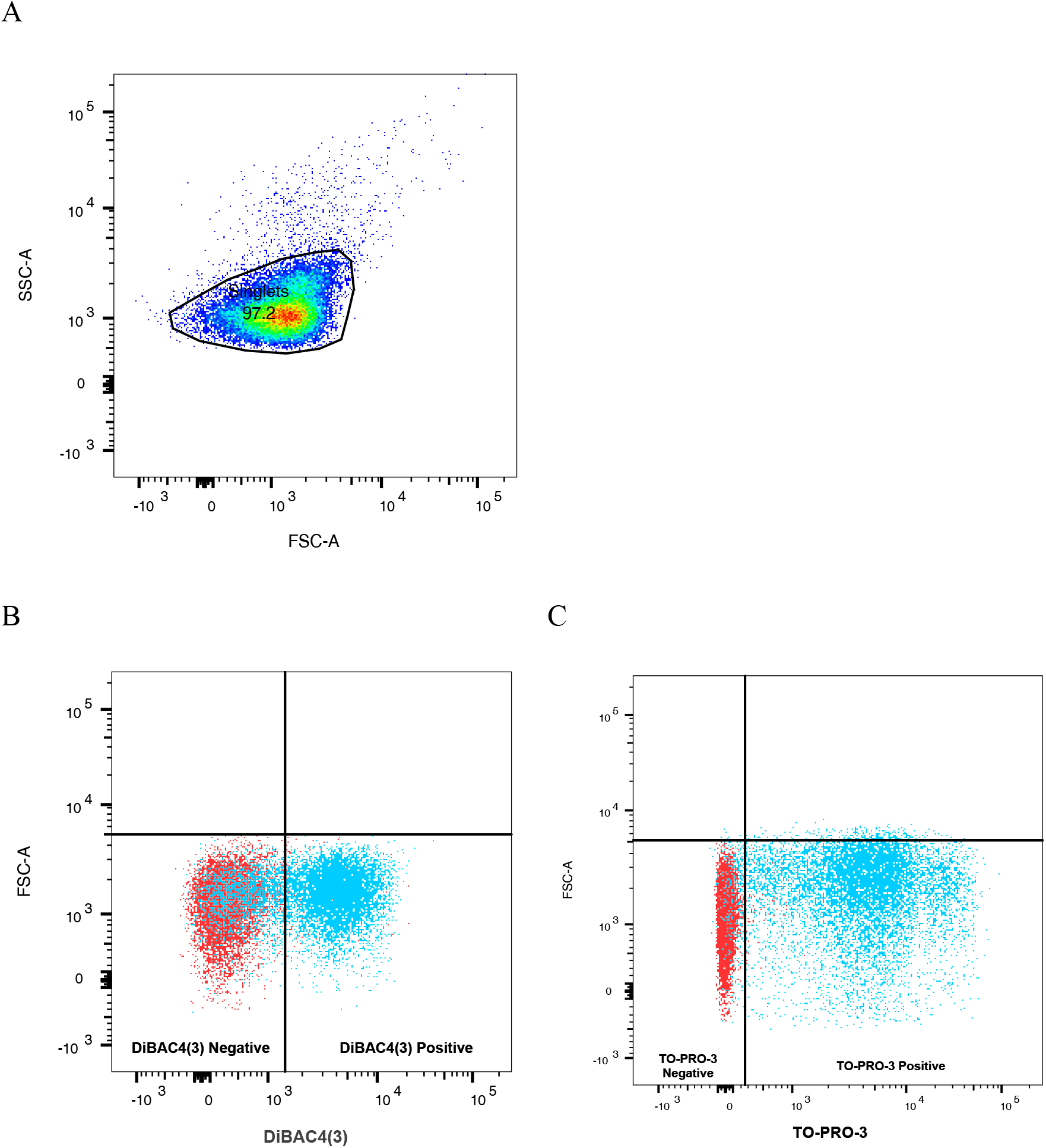
Representative flow data shown. Mid-exponential *P. aeruginosa* PAO1-LAC-EcPore cells were either untreated or treated with CCCP (100 µM) for 2 hours. Cells were stained with DiBAC4(3) to determine the number of cells with depolarised membranes. A) Scatter plot of side scatter area (SSC-A) versus forward scatter area (FSC-A), singlets gate excludes debris. B) Scatter plot of FSC-A versus DiBAC_4_(3) fluorescence measured in the FITC-A channel. The lower left quadrant represents cells negative for DiBAC_4_(3) fluorescence and the lower right quadrant represents cells positive for DiBAC_4_(3) fluorescence. C) Scatter plot of FSC-A versus TO-PRO^®^-3 fluorescence measured in the APC-A channel. The lower left quadrant represents cells negative for TO-PRO^®^-3 fluorescence and the lower right quadrant represents cells positive for TO-PRO^®^-3 fluorescence.

